# Species distribution modeling and conservation assessment of the black-headed night monkey (*Aotus nigriceps*) – A species of Least Concern that faces widespread anthropogenic threats

**DOI:** 10.1101/2020.05.20.107383

**Authors:** William D. Helenbrook, Jose W. Valdez

## Abstract

Deforestation rates in the Brazilian Amazon have been steadily increasing since 2007. Recent government policy, projected growth of agriculture, and expansion of the cattle industry is expected to further pressure primates within the Amazon basin. In this study, we examined the anthropogenic impact on the widely distributed black-headed night monkey, *Aotus nigriceps*, whose distribution and population status have yet to be assessed. We 1) modeled species distribution in *A. nigriceps*; 2) estimated impact of habitat loss on population trends; and 3) highlight landscape-based conservation actions which maximize potential for their long-term sustainability. We found the black-headed night monkey to be restricted by several biotic and environmental factors including forest cover, elevation, isothermality, and precipitation. Over the last two decades, over 132,908 km^2^ of tree cover (18%) has been lost within their documented range. We found this species occupies only 49% of habitat within in their range, a loss of 19% from their estimated 2000 distribution, and just over 34% of occupied areas were in protected areas. Projected deforestation rates of *A. nigriceps* equates to an additional loss of 23,084 km^2^ of occupied habitat over the next decade. This study suggests that although classified as a species of Least Concern, *A. nigriceps* may have a much smaller range and is likely more at risk than previously described. The future impact of continued expansion of mono-cultured crops, cattle ranching, and wildfires is still unknown. However, expanded use of participatory REDD+, sustainable agroforestry in buffer zones, secured land tenor for indigenous communities, wildlife corridors, and the expansion of protected areas can help ensure viability for this nocturnal primate and other sympatric species throughout the Amazon Basin.

## Introduction

Deforestation rates in the Amazon rainforest are at their highest levels since 2007 due to expansion of mono-cultured agriculture (e.g., palm oil and soy), cattle ranching, wildfires, and urban expansion (Davidson et al., 2012; PRODES 2020); with models predicting up to 40% of all Amazonian forests will be lost by 2050 (Gomes et al., 2019). These at-risk areas overlap with many neotropical primate species whose populations may suffer large declines. This includes the black-headed night monkey, *Aotus nigriceps* (Dollman 1909), one of eleven currently recognized species of night monkey found in Central and South America (Hershkovitz 1983; Defler and Bueno 2007; Plautz et al. 2009; Babb et al. 2011; Ruiz-Garcia et al., 2011).

*Aotus nigriceps* is a wide-ranging nocturnal monkey species found across much of the central and upper Amazon, including degraded forest, agroforested landscapes, and areas in close proximity to humans (Helenbrook et al., 2020). Nevertheless, while night monkeys are adaptable and persist in degraded landscapes, they still require connective forest cover, access to seasonally available fruiting trees, sleeping sites, and are restricted by environmental conditions (Aquino & Encarnación 1994; Helenbrook et al., 2020). Although thought to be a common species, it is currently listed as Appendix II of CITES, and the impact of anthropogenic disturbances on the long-term viablity of *A. nigriceps* remains unknown.

*Aotus nigriceps* is found in Southern Peru, northern Bolivia, and central-western Brazil. However, discrepancies remain regarding the extent of their eastern distribution in the state of Rondonia and additional taxonomic analysis is needed from the area (e.g., Pieczarka et al. 1993; Plautz et al. 2009; Menezes et al. 2010; Babb et al. 2011; Ruiz-Garcia et al., 2011). The species is not considered under threat due to an extensive range, suspected large and sustained population size, presumed low risk of extinction throughout the species range (Shanee et al. 2018), and high density estimates ranging from 19-50 individuals/km^2^ (Helenbrook et al., 2020). However, no known population size estimates or trends have been described. This is concerning since large portions of its range overlap with areas experiencing some of the highest deforestation rates in the world (Estrada et al., 2018). Besides habitat loss, their long-term viability is also projected to be further impacted by altered phenology and increased average maximum temperatures from climate change, which will force species to adapt their distribution or face increased risk from extinction (e.g., Raghunathan et al., 2014; Cohn et al., 2019; Carvalho et al., 2019a).

Assessing the population status of *Aotus nigriceps*, and any species in general, requires accurately describing their distribution, baseline population size, and predicting future population growth or loss. This information can be used to build predictive models that can indicate priority regions for conservation action. Species distribution modeling is a well-known approach, which predicts a species distribution across a landscape based on the environmental conditions necessary for species occupancy (e.g., Bett et al., 2012; Kamilar & Tecot 2015; Rabelo et al., 2018). The MaxEnt program is especially suited for this task and is considered the most accurate maximum entropy machine learning model, achieving high predictive accuracy without true absence data, incomplete data, and small sample sizes (Phillips & Dudik 2008; Merow et al. 2013). Maxent has recently been used in primate studies to determine current and future distribution of snub-nosed monkeys (Wong et al. 2013; Luo et al. 2014; Nüchel et al., 2018), habitat suitability of Sichuan golden monkeys (e.g., Liu et al. 2017), potential distribution of Caatinga howler monkeys (Filho and Palmeirim 2019), and conservation efforts for the brown-headed spider monkey (Peck et al. 2011). However, it is typically not used for species of Least Concern and only after a species is considered under threat.

For this study, we used species distribution modeling with MaxEnt to estimate *A. nigriceps* distribution and then projected density measurements across their predicted distribution. The objectives of this study are to: 1) develop baseline distribution models on the occupancy of *A. nigriceps* utilizing ecological, biogeographical, and environmental data; 2) project population losses due to habitat loss and degradation; and 3) identify priority conservation actions needed to sustain black-headed night monkey populations.

## Methods

### Species occurrence and environmental data

We constructed an occurrence data set (N=75) of *Aotus nigriceps* observations throughout their range (10.13140/RG.2.2.33456.58886) (Fig. 1). We used several available sources, including published manuscripts (Wright 1996; Helenbrook et al. 2020) and field records provided by the Wildlife Conservation Society and iNaturalist (Lopez-Strauss et al., 2016) (Fig. 2a). To ensure accuracy, we only included data which had confirmed species identity and accurate geographic location. All points were confirmed to ensure coordinates aligned with described distributions and associated habitat. All records were then plotted in QGIS (3.12.1) and created a polygon layer that included all records and followed major river barriers. The polygon represented the maximum species’ extent of occurrence described by several peer-reviewed sources (e.g., Plautz et al. 2009; Babb et al. 2011).

**FIG. 1:**
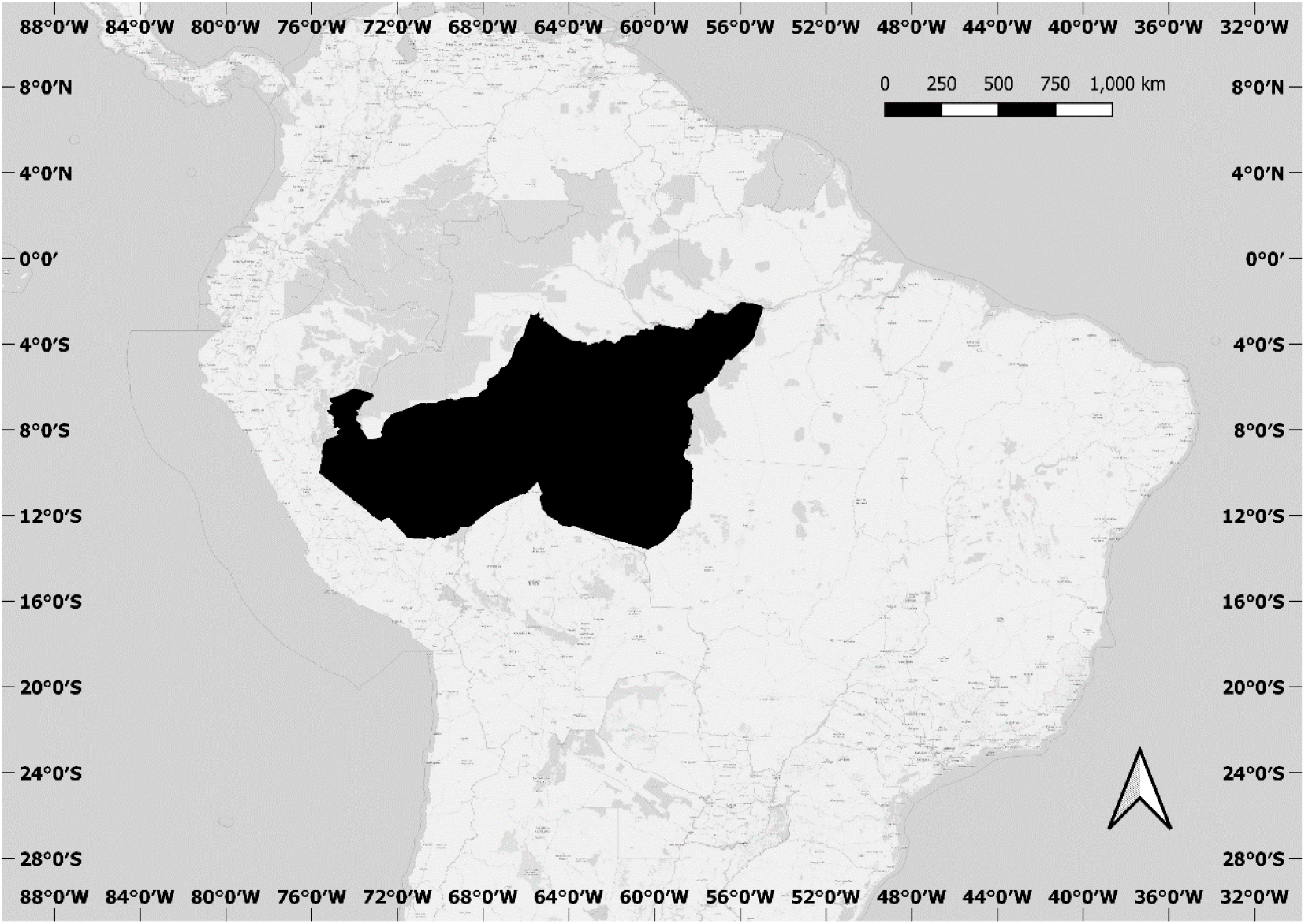
Currently described *Aotus nigriceps* range based on biogeographical limitations and literature.

**FIG. 2:**
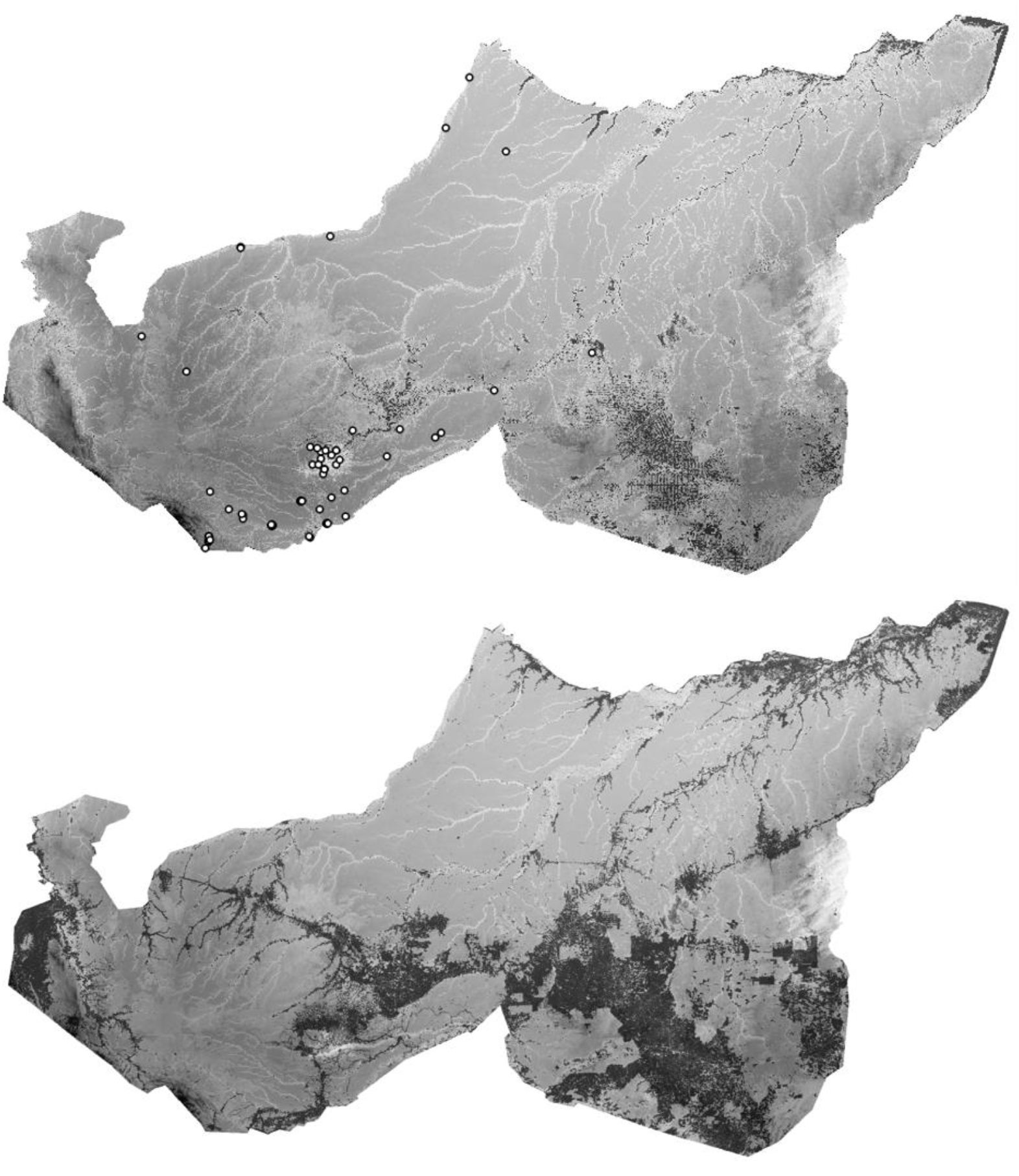
a) *Aotus nigriceps* predicted distribution based on the maximum test sensitivity plus specificity threshold in a) 2000 and b) 2018. Darker areas represent lower probability. a) Includes presence data used in Maxent models.

We used bioclimatic variables as environmental predictors based on their presumed ecological or physiological relationship to black-headed night monkey suitability as previously used in other neotropical primate studies (e.g., Booth et al., 2014; Holzmann et al., 2014; Rabelo et al., 2018). Climatic variables were downloaded from WorldClim dataset and included annual mean temperature, annual temperature range, annual precipitation, and isothermality (defined as mean diurnal range [mean of monthly (max temperature – min temp)] divided by temperature annual range) (Fick & Hijmans 2017). For elevation, we used the Shuttle Radar Topographic Mission elevation dataset (SRTM 2020). We also obtained percent forest cover data from 2000, forest gain from 2000-2012, and forest loss between 2000-2018 from Earth Engine Partners (Hansen et al. 2013). Autocorrelated variables greater than 0.7 were removed from the analyses to avoid overfitting. To calculate the most current cover percentage for the year 2018, the cumulative gain and loss between 2000 and 2018 were added and subtracted from the 2000 cover layer within their range, respectively. We calculated protected area within *A. nigriceps* range and predicted distribution using polygons obtained from Protected Planet (UNEP-WCMC & IUCN 2020). All layers utilized in the study were in 30 seconds (1 km) spatial resolution.

### Distribution models

We used MaxEnt (Version: 3.4.1) to develop a correlative species distribution model for *Aotus nigriceps*, relating known geographic occurrences to available environmental variables to estimate suitable habitat (Phillips & Dudík, 2008; Franklin 2013). To model their predicted distribution, we used 2000 and 2018 forest cover layers to predict their most current distribution. We used 70% of known species’ localities as training data and 30% of the localities were set aside as test data. A bootstrap method was used with a maximum of 5,000 iterations, 10,000 randomly selected background points, a prevalence of 0.75, and 15 replicates with a random seed. Since there was a spatial sampling bias with many of the observations, a kernel density analysis was conducted using QGIS (3.12.1), and this raster layer representing the intensity of points was then used as a bias file in Maxent (Brown 2014). A regularization multiplier of 3 was also added to further constrain the model and avoid overfitting, reduce complexity, and generate less localized predictions (Anderson & Gonzalez, 2011; Merow et al., 2013; Shcheglovitova & Anderson, 2013; Morales et al., 2017).

To assess model discriminatory power and compare performance of species distribution models, we measured the area under the receiver operating characteristic curve (AUC) (Fielding & Bell, 1997; Babar et al., 2012; Phillips & Dudík, 2008; Stohlgren et al., 2011). To measure variable importance and determine which variables contributed most to the model, we performed a Jackknife analysis with and without each variable in isolation. Habitat was considered suitable enough to be occupied based on the maximum specificity and sensitivity threshold which has been found to produce the most accurate results (Liu et al. 2005, Jimenez-Valverde and Lobo 2007). We then calculated total area of suitable habitat using QGIS.

### Population Size Estimates and Threat Assessment

To estimate loss of habitat over time we 1) calculated tree cover loss within their documented range from Global Forest Watch data layers (Hansen et. al. 2013), and 2) calculated the area of their occurrence between 2000 and 2018 using the MaxEnt modeled species distribution. We then used upper and lower population density estimates obtained from Helenbrook et al. (2020) to extrapolate across projected habitat using area of occurrence calculations. Future deforestation projections in the Brazilian Amazon were based on rates calculated by Butler (2020). We highlight management practices needed to minimize habitat loss, maintain connectivity, and restore populations regionally.

## Results

### Species Distribution Modeling

MaxEnt models showed fair support for predicting observed distribution (AUC = 0.688), representative of generalized species. Mean temperature and annual temperature range were removed from the final models due to high correlation with other variables. Occupancy increased with greater forest cover (range 0 - 100%) and precipitation (range 819 – 4,797mm), decreased with isothermality (range 6.4 - 8.7%) and elevation (range 1 - 3,062 m) (Fig. 3a). Cover and isothermality had the highest permutation importance (Table 1). Cover and elevation were the most effective variables for predicting the distribution of the occurrence data set aside for training, cover for testing, and cover, isotherm, and precipitation when predictive performance was measured using AUC (Fig. 3b). The largest contributing environmental variable when omitted in all jackknife tests was elevation, isothermality, and precipitation, suggesting that they contribute the most information not found in the other variables.

**TABLE 1:** Relative contribution estimates of environmental variables to the MaxEnt model predicting *Aotus nigriceps* occupancy. For percent contribution, the increase in regularized gain for each iteration of the training algorithm is added to the contribution of the corresponding variable or subtracted if the change to the absolute value of lambda is negative. For permutation importance, the values of the environmental variable on training presence and background data are randomly permuted. The model is reevaluated on the permuted data, and the resulting drop in training AUC is normalized to percentages. Values shown are averages over replicate runs.

| Variable | Percent contribution | Permutation importance |
| --- | --- | --- |
| Forest cover | 58.9 | 59.6 |
| Elevation | 23.5 | 1.7 |
| Isothermality | 16.4 | 32.8 |
| Precipitation range | 1.9 | 5.9 |

**FIG. 3a.**
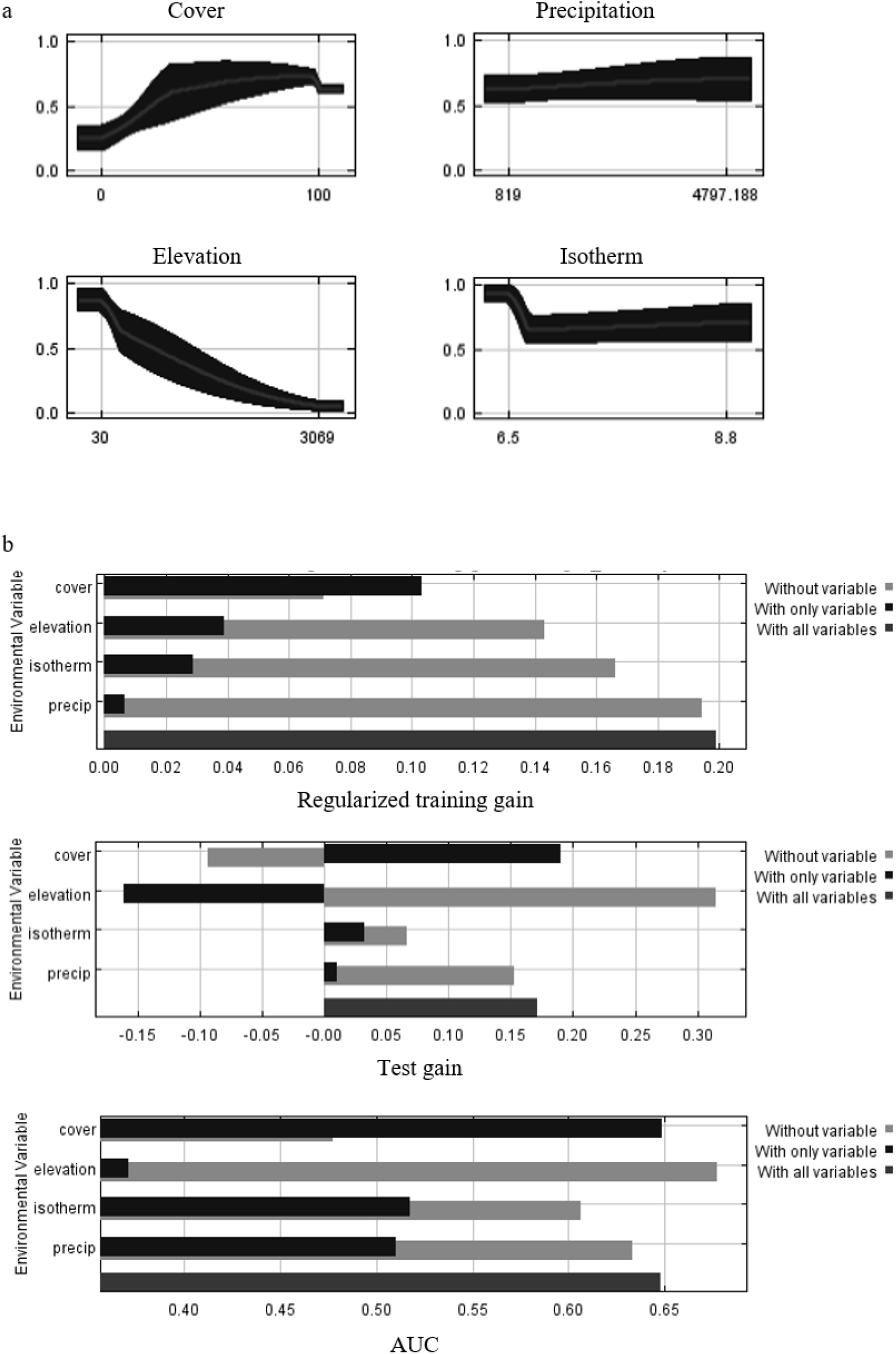
The predicted probabilities of *Aotus nigriceps* occupancy of each environmental variable, keeping all other variables at their average sample. The curves represent the mean response of 15 replicate MaxEnt runs (red) and +/- one standard deviation (blue). FIG. 3b. Jackknife tests for variable importance for regularized training gain, test gain, and AUC averaged over 15 replicate bootstrap runs for predicting *Aotus nigriceps* distribution modeling.

The estimated current extent of *Aotus nigriceps* was 1,591,711 km^2^ based on the available literature (Hershkovitz 1983; Defler and Bueno 2007; Plautz et al. 2009; Babb et al. 2011; Ruiz-Garcia et al., 2011: Fig. 1). We calculated a loss of 18% of forest cover between 2000-2018 satellite imagery data (Fig 4a). Protected areas represented 14% of forest cover within their range in 2000 and exhibited forest cover loss of 5% over the past decade. Their predicted distribution based on the maximum test sensitivity plus specificity threshold indicated they occupied 49% (779,938 km^2^) of their known range (Fig. 2b), a loss of 19% from the predicted 60% in 2000 (Fig. 2a). Of their predicted distribution only a third (34%) was in protected areas (Fig. 4b).

**FIG 4:**
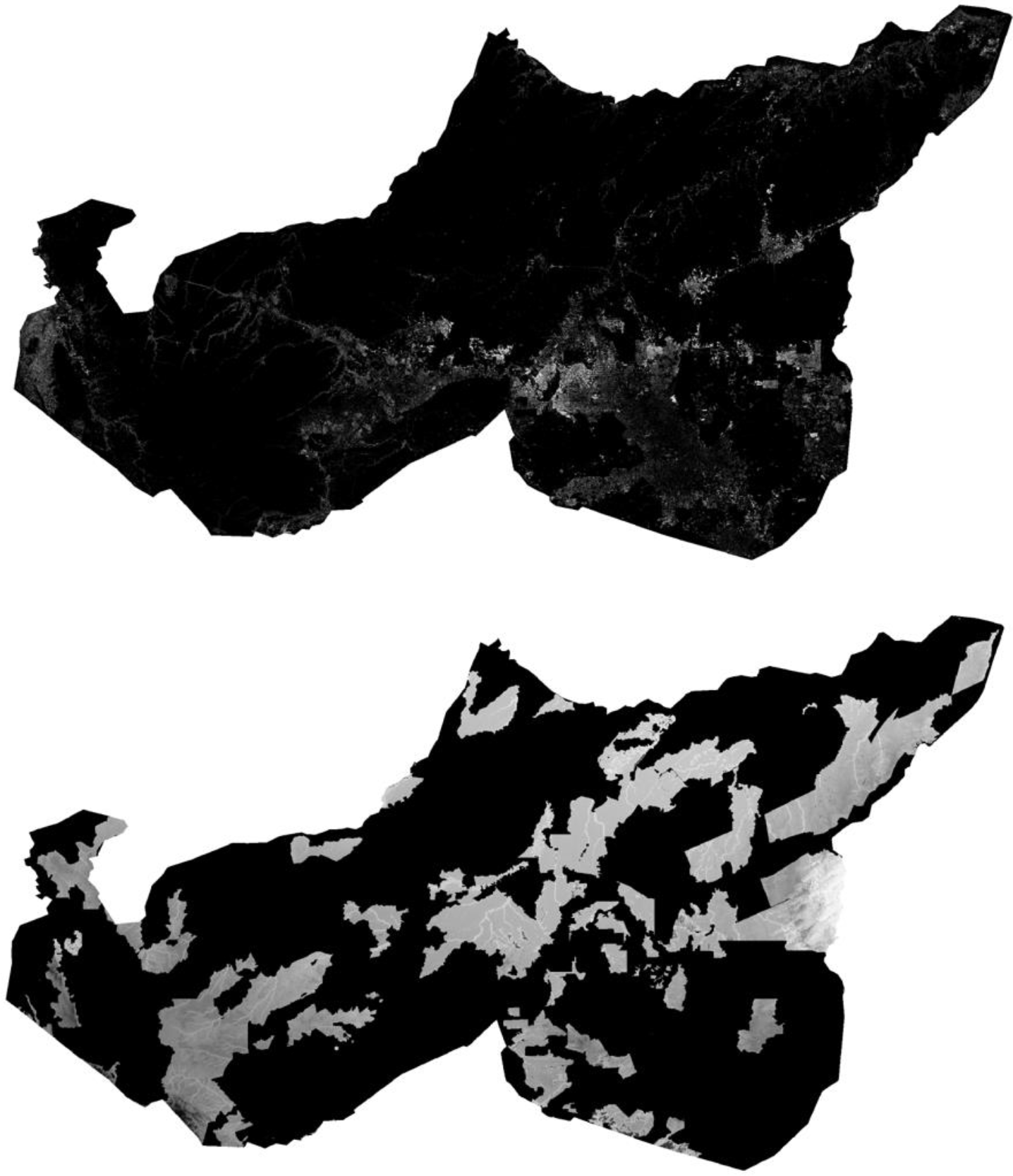
a) Forest loss between 2000-2018 based on satellite imagery obtained from Earth Engine Partners. We calculated a loss of 18% of forest cover between 2000-2018 satellite imagery data b) Protected areas in gray. Lighter gray areas have higher probability of occupancy.

Current population estimates using modeled viable habitat coupled with density estimates from previous studies (i.e., 19-50 individuals per km^2^) range from 14,818,822 to 38,996,900 individuals (Helenbrook et al., 2020). Based on contraction of suitable habitat between 2000-2018, we estimate a loss of 3,326,672-8,754,400 individuals. Current extrapolated deforestation rates across present-day modeled distribution of *A. nigriceps* equates to an additional loss of 23,084 km^2^ of habitat over the next decade.

## Discussion

This is the first study to assess the species distribution, population status, and conservation management implications for the black-headed night monkey. Although considered Least Concern due to their adaptability to a variety of habitat types including the moist forests of the Southwest Amazon, Jurua-Purus, Purus-Maderia, and Maderia-Tapajos, *A. nigriceps* distribution appears restricted by biotic and environmental factors. In particular, they were more likely to occupy habitats represented by low altitudes, low isothermality, higher precipitation, and high forest cover. We found that although not considered a species of concern due to its broad distribution across several Amazon Basin ecoregions, they currently occupy a much smaller distribution than the previously described extent of their range would suggest. Even two decades ago, their distribution was only half of their presumed extent of occurrence. Furthermore, we estimate that between 2000 and 2018, this species has been reduced by at least 3 million individuals. Although this species is currently considered common, they will likely continue to decline significantly due to widespread anthropogenic pressures, especially since less than a quarter of their suitable habitat is within protected areas.

This study provides evidence that *A. nigriceps* has experienced sharp declines in forest cover in portions of their range over the last couple of decades and will continue to experience habitat loss and degradation resulting in population decline over the coming decade. Deforestation rates have surged recently in Brazil driven by heavy pressure from cattle ranching and mono-cultured agriculture (Gomes et al., 2019; IMAZON 2020). Some of the highest deforestation rates in the Amazon occur in the Brazilian States of Pará, Rondonia, and Mato Grosso where this nocturnal species occurs. Systematic conservation planning based solely on the black-headed night monkey appears premature at this time due to the broad distribution of the species and without a thorough analysis of underlying taxonomy and molecular variability throughout its range. However, the expansion of several existing conservation strategies would benefit *A. nigriceps* and contribute to overall biodiversity conservation.

Although extensive protected areas and indigenous territories exist in Peru, Bolivia, and Brazil, though level of protection varies widely (Amazonia Socioambiental 2019; Paiva et al. 2020). Forest loss and degradation still occurs within protected and indigenous lands, though the degree to which these areas is impacted depends on the type of protected area and management (e.g., indigenous, federal, or state: Herrera et al., 2019). Regardless of governance, primates clearly benefit from protected areas (e.g., Paim et al. 2018). Continued support for participatory and jurisdictional REDD+ programs also appears effective though this approach requires realistic expectations among local communities regarding benefits and participation, secured land tenor, and a mix of intervention strategies including incentives, disincentives, and introduction of alternative livelihoods (e.g., Davis and Goldman 2017; Simonet et al., 2019; Dupuits & Cronkleton 2020). Agroforestry research related to night monkey species is limited, though a study in the northern Andes suggests that coffee plantations can benefit night monkeys while also providing farmers with income (Guzmán et al., 2016). We have previously documented that the black-headed night monkey is quite common in secondary forest and that they make use of orchards (Helenbrook et al., 2020). The main issue would appear to be whether connected canopy is maintained in order for night monkeys to traverse a complex matrix with forest fragmentation and diverse land uses.

Here, we presented an assessment of the current state of forest degradation and loss in the Amazon, coupled with an analysis of the associated anthropogenic impact on a common primate species. Our goal is not to suggest that the black-headed night monkey is under threat of extinction but to show that even widely distributed primates face extensive risks. More specialized sympatric primate species likely face a far greater threat of habitat loss and plummeting population size. By using the black-headed night monkey species as an indicator of landscape and forest health, solutions that mitigate their decline would also favor those that are much more vulnerable – either because they are more sensitive to habitat disturbances, have far smaller numbers, or have yet to even be discovered. There is also the issue of *A. nigriceps* taxonomic uncertainty. There may very well be undescribed conservation management units across this widely distributed species – or currently unrecognized subspecies driven by riverine barriers. This species is distributed across the Amazon but could potentially include units that are each under their own unique stresses and face reduced viability compared to the whole. The range of *A. nigriceps* is also likely to change with continued molecular and morphological analysis. Nevertheless, this study provides critical information of areas highly likely to have black-headed monkeys which can be used for future conservation monitoring and planning. This species has experienced extensive habitat loss over the past couple of decades and there will likely be continued habitat loss and degradation, threatening this relatively common species. A regional landscape-level conservation plan that maintains forest connectivity and minimizes habitat loss would benefit not only the black-headed night monkey but the many other sympatric species who are under increasing threat in the region.

## Data Availability

The data that support the findings of this study are openly available in ResearchGate: DOI: 10.13140/RG.2.2.33456.58886.

## Author contributions

Study design, data collection, analysis, and writing: WDH; Modeling and assistance with writing: JWV

## Acknowledgements

Thank you to Hillary Fenrich, Sheridan Plummer, and Ben Sharaf who assisted in original model development. Thank you to Rob Wallace for sharing Wildlife Conservation Society occurrence data from Bolivia. We are indebted to the anonymous reviewers who provided constructive feedback.

## Ethical Standards

This research complied with the American Journal of Primatology Code of Conduct.

## References

Amazonia Socioambiental (2019). Amazonia Network of Georeferenced Socio-Environmental Information. https://www.amazoniasocioambiental.org/en/maps/ [accessed 7 May 2020].

Anderson, R.P. & Gonzalez, I. (2011) Species-specific tuning increases robustness to sampling bias in models of species distributions: an implementation with MaxEnt. Ecological Modelling, 222, 2796–2811. DOI: 10.1016/j.ecolmodel.2011.04.011

Aquino, R., & Encamación, F. (1994). Owl monkey populations in Latin America: field work and conservation. Aotus: The Owl Monkey, 59–95.

Babar, S., Amarnath, G., Reddy, C.S., Jentsch, A., & Sudhakar, S. (2012). Species distribution models: ecological explanation and prediction of an endemic and endangered plant species (Pterocarpus santalinus Lf). Current Science, 1157–1165.

Bett, N.N., Blair, M.E., & Sterling, E.J. (2012). Ecological niche conservatism in doucs (Genus Pygathrix). International Journal of Primatology, 33, 972–988. DOI: 10.1007/s10764-012-9622-3

Brown, J. L. (2014). SDM toolbox: a python-based GIS toolkit for landscape genetic, biogeographic and species distribution model analyses. Methods in Ecology and Evolution, 5, 694–700. DOI: 10.1111/2041-210X.12200

Butler, R. 2020. What’s the deforestation rate in the Amazon? https://rainforests.mongabay.com/amazon/deforestation-rate.html [accessed 8 May 2020].

Carvalho, J.S., Graham, B., Rebelo, H., Bocksberger, G., Meyer, C.F., Wich, S., et al. (2019a). A global risk assessment of primates under climate and land use/cover scenarios. Global Change Biology, 25, 3163–3178. DOI: 10.1111/gcb.14671

Carvalho, W.D., Mustin, K., Hilário, R.R., Vasconcelos, I.M., Eilers, V., & Fearnside, P.M. (2019b). Deforestation control in the Brazilian Amazon: A conservation struggle being lost as agreements and regulations are subverted and bypassed. Perspectives in Ecology and Conservation, 17, 122–130. DOI: 10.1016/j.pecon.2019.06.002

Cohn, A.S., Bhattarai, N., Campolo, J., Crompton, O., Dralle, D., Duncan, J., & Thompson, S. (2019). Forest loss in Brazil increases maximum temperatures within 50 km. Environmental Research Letters, 14, 084047. DOI: 10.1088/1748-9326/ab31fb

Davidson, E.A., de Araújo, A.C., Artaxo, P., Balch, J.K., Brown, I.F., Bustamante, M.M., et al. (2012). The Amazon basin in transition. Nature, 481, 321. DOI: 10.1038/nature10717

Davis, A., & Goldman, M.J. (2019). Beyond payments for ecosystem services: considerations of trust, livelihoods and tenure security in community-based conservation projects. Oryx, 53, 491–496. DOI: 10.1017/S0030605317000898

Defler, T.R., & Bueno, M.L. (2007). Aotus diversity and the species problem. Primate Conservation, 22, 55–70. DOI: 10.1896/052.022.0104

Dupuits, E., & Cronkleton, P. (2020). Indigenous tenure security and local participation in climate mitigation programs: Exploring the institutional gaps of REDD+ implementation in the Peruvian Amazon. Environmental Policy and Governance, 1–12. DOI: 10.1002/eet.1888

Estrada, A., Raboy, B.E., & Oliveira, L.C. (2012). Agroecosystems and primate conservation in the tropics: a review. American Journal of Primatology, 74, 696–711. DOI: 10.1002/ajp.22033

Estrada, A., Garber, P.A., Mittermeier, R.A., Wich, S., Gouveia, S., Dobrovolski, R., et al. (2018). Primates in peril: the significance of Brazil, Madagascar, Indonesia and the Democratic Republic of the Congo for global primate conservation. PeerJ, 6, e4869. DOI: 10.7717/peerj.4869

Fick, S.E. & Hijmans, R.J. (2017). WorldClim 2: new 1km spatial resolution climate surfaces for global land areas. International Journal of Climatology, 37, 4302–4315. DOI: 10.1002/joc.5086

Fielding, A.H., & Bell, J.F. (1997). A review of methods for the assessment of prediction errors in conservation presence/absence models. Environmental Conservation, 24, 38–49. DOI: 10.1017/S0376892997000088

Filho, R., & Palmeirim, J. M. (2019). Potential distribution of and priority conservation areas for the Endangered Caatinga howler monkey Alouatta ululata in north-eastern Brazil. Oryx, 1–9. DOI: 10.1017/S0030605318001084

Franklin, J. (2013). Species distribution models in conservation biogeography: developments and challenges. Diversity and Distributions, 19, 1217–1223. DOI: 10.1111/ddi.12125

Freire Filho, R., & Palmeirim, J. M. (2019). Potential distribution of and priority conservation areas for the Endangered Caatinga howler monkey Alouatta ululata in north-eastern Brazil. Oryx, 1–9. DOI: 10.1017/S0030605318001084

Gomes, V.H., Vieira, I.C., Salomão, R.P., & Steege, H. (2019). Amazonian tree species threatened by deforestation and climate change. Nature Climate Change, 9, 547–553. DOI: 10.1038/s41558-019-0500-2

Guzmán, A., Link, A., Castillo, J.A., & Botero, J.E. (2016). Agroecosystems and primate conservation: Shade coffee as potential habitat for the conservation of Andean night monkeys in the northern Andes. Agriculture, Ecosystems & Environment, 215, 57–67. DOI: 10.1016/j.agee.2015.09.002

Hansen, M.C., Potapov, P.V., Moore, R., Hancher, M., Turubanova, S.A. Tyukavina, A. D. et al. (2013). High-resolution global maps of 21st-century forest cover change. Science, 342, 850–853. DOI: 10.1126/science.1244693

Helenbrook, W.D. & Valdez, J.W. (2020). Species distribution modeling. ResearchGate: DOI 10.13140/RG.2.2.33456.58886.

Helenbrook, W.D., Wilkinson, M.L., & Suarez, J.A. (2020). Habitat use, fruit consumption, and population density of the black-headed night monkey, Aotus nigriceps, in Southeastern Peru. Acta Amazonica, 50, 37–43. DOI: 10.1590/1809-4392201900172

Herrera, D., Pfaff, A., & Robalino, J. (2019). Impacts of protected areas vary with the level of government: Comparing avoided deforestation across agencies in the Brazilian Amazon. Proceedings of the National Academy of Sciences, 116, 14916–14925. DOI: 10.1073/pnas.1802877116

Hershkovitz, P. (1983). Two new species of night monkeys, genus Aotus (Cebidae, Platyrrhini): a preliminary report on Aotus taxonomy. American Journal of Primatology, 4, 209–243. 10.1002/ajp.1350040302

Holzmann, I., Agostini, I., DeMatteo, K., Areta, J.I., Merino, M.L., & Di Bitetti, M.S. (2014). Using species distribution modeling to assess factors that determine the distribution of two parapatric howlers (Alouatta spp.) in South America. International Journal of Primatology, 36, 18-32. 10.1007/s10764-014-9805-1

IMAZON 2020. Instituto do Homem e Meio Ambienta da Amazonia. https://imazon.org.br/publicacoes/boletim-do-desmatamento-da-amazonia-legal-janeiro-2019-sad/ [accessed 29 April 2020].

Ingberman, B., Fusco-Costa, R., & Monteiro-Filho, E. L. (2016). A current perspective on the historical geographic distribution of the endangered muriquis (Brachyteles spp.): Implications for conservation. PloS One, 11, e0150906. DOI: 10.1371/journal.pone.0150906

Jiménez-Valverde, A., & Lobo, J. M. (2007). Threshold criteria for conversion of probability of species presence to either–or presence–absence. Acta oecologica, 31, 361–369. DOI: 10.1016/j.actao.2007.02.001

Kamilar, J.M., & Tecot, S.R. (2016). Anthropogenic and climatic effects on the distribution of Eulemur species: an ecological niche modeling approach. International Journal of Primatology, 37, 47–68. DOI: 10.1007/s10764-015-9875-8

Liu, C., Berry, P. M., Dawson, T. P., & Pearson, R. G. (2005). Selecting thresholds of occurrence in the prediction of species distributions. Ecography, 28, 385–393. DOI: 10.1111/j.0906-7590.2005.03957.x

Lopez-Strauss, H., R.B. Wallace, N. Mercado & Z.R. Porcel. (2016). Current knowledge on primate distribution and conservation in Bolivia. Chapter 19 pp. 575-640. In: Phylogeny, Molecular Population Genetics, Evolutionary Biology and Conservation of the Neotropical Primates. Eds: Ruiz Garcia, M. & J. Shostell. Nova Science Publisher, New York.

Luo, Z., Zhou, S., Yu, W., Yu, H., Yang, J., et al. (2015). Impacts of climate change on the distribution of Sichuan snub-nosed monkeys (Rhinopithecus roxellana) in Shennongjia area, China. American Journal of Primatology, 77, 135–151. DOI: 10.1002/ajp.22317

Merow, C., Matthew J.S., & Silander, JA. (2013). A practical guide to MaxEnt for modeling species’ distributions: what it does, and why inputs and settings matter. Ecography, 36, 1058-1069. 10.1111/j.1600-0587.2013.07872.

Meyer, A. L., Pie, M. R., & Passos, F. C. (2014). Assessing the exposure of lion tamarins (Leontopithecus spp.) to future climate change. American Journal of Primatology, 76, 551–562. DOI: 10.1002/ajp.22247

Morales N.S., Fernández, I.C., & Baca-González, V. (2017). MaxEnt’s parameter configuration and small samples: are we paying attention to recommendations? A systematic review. PeerJ. 5, e3093. DOI: 10.7717/peerj.3093

Nüchel, J., Bøcher, P. K., Xiao, W., Zhu, A. X., & Svenning, J. C. (2018). Snub-nosed monkeys (Rhinopithecus): potential distribution and its implication for conservation. Biodiversity and Conservation, 27, 1517–1538. DOI: 10.1007/s10531-018-1507-0

Paim, F.P., El Bizri, H.R., Paglia, A.P., & Queiroz, H.L. (2019). Long-term population monitoring of the threatened and endemic black-headed squirrel monkey (Saimiri vanzolinii) shows the importance of protected areas for primate conservation in Amazonia. American Journal of Primatology, 81, e22988. DOI: 10.1002/ajp.22988

Paiva, P.F., Ruivo, M.D., da Silva Júnior, O.M., Maciel, M.D., Braga, T. G., de Andrade, M.M. et al. (2020). Deforestation in protect areas in the Amazon: a threat to biodiversity. Biodiversity and Conservation, 29, 19–38. DOI: 10.1007/s10531-019-01867-9

Peck, M., Thorn, J., Mariscal, A., Baird, A., Tirira, D., & Kniveton, D. (2011). Focusing conservation efforts for the critically endangered brown-headed spider monkey (Ateles fusciceps) using remote sensing, modeling, and playback survey methods. International Journal of Primatology, 32, 134–148. DOI: 10.1007/s10764-010-9445-z

Peres, C.A. (1993). Structure and spatial organization of an Amazonian terra firme forest primate community. Journal of Tropical Ecology, 259–276. DOI: 10.1017/S026646740000729X

Peres, C. A. (1997). Primate community structure at twenty western Amazonian flooded and unflooded forests. Journal of Tropical Ecology, 381–405. DOI: 10.1017/S0266467400010580

Peterson, A.T., & Soberón, J. (2012). Species distribution modeling and ecological niche modeling: getting the concepts right. Natureza & Conservação, 10, 102–107. DOI: 10.4322/natcon.2012.019

Phillips, S. J., & Dudík, M. (2008). Modeling of species distributions with MaxEnt: new extensions and a comprehensive evaluation. Ecography, 31, 161–175. DOI: 10.1111/j.0906-7590.2008.5203.x

PRODES (2020). Prodes deforestation. Global Forest Watch. http://data.globalforestwatch.org/datasets/prodes-deforestation [accessed 8 May 2020].

Rabelo, R.M., Gonçalves, J.R., Silva, F.E., Rocha, D.G., Canale, G.R., Bernardo, C.S., et al. (2018). Predicted distribution and habitat loss for the endangered black-faced black spider monkey Ateles chamek in the Amazon. Oryx, 1–7. DOI: 10.1017/S0030605318000522

Raghunathan, N., François, L., Huynen, M.C., Oliveira, L.C., & Hambuckers, A. (2015). Modelling the distribution of key tree species used by lion tamarins in the Brazilian Atlantic forest under a scenario of future climate change. Regional Environmental Change, 15, 683–693. DOI: 10.1007/s10113-014-0625-9

Ribas, C.C., Aleixo, A., Nogueira, A.C., Miyaki, C.Y., & Cracraft, J. (2012). A palaeobiogeographic model for biotic diversification within Amazonia over the past three million years. Proceedings of the Royal Society B: Biological Sciences, 279, 681–689. DOI: 10.1098/rspb.2011.1120

Ruiz-García, M., Vásquez, C., Camargo, E., Leguizamón, N., Gálvez, H. Vallejo, et al. (2011). Molecular phylogenetics of Aotus (Platyrrhini, Cebidae). International Journal of Primatology, 32, 1218. DOI: 10.1007/s10764-011-9539-2

Sarma, K., Kumar, A., Krishna, M., Medhi, M., & Tripathi, O. P. (2015). Predicting suitable habitats for the vulnerable Eastern Hoolock Gibbon, Hoolock leuconedys, in India using the MaxEnt Model. Folia Primatologica, 86, 387–397. DOI: 10.1159/000381952

Shanee, S., Alves, S.L., Calouro, A.M., Lynch Alfaro, J., Messias, M. (2018). Aotus nigriceps. The IUCN Red List of Threatened Species e.T41542A17923573.

Shcheglovitova, M. & Anderson, R.P. (2013) Estimating optimal complexity for ecological niche models: A jackknife approach for species with small sample sizes. Ecological Modelling, 269, 9–17. DOI: 10.1016/j.ecolmodel.2013.08.011

Silva, S.M., Peterson, A.T., Carneiro, L., Burlamaqui, T.C., Ribas, C.C., Sousa-Neves, T., et al. (2019). A dynamic continental moisture gradient drove Amazonian bird diversification. Science Advances, 5, eaat5752. DOI: 10.1126/sciadv.aat5752

Simonet, G., Subervie, J., Ezzine-de-Blas, D., Cromberg, M., & Duchelle, A.E. (2019). Effectiveness of a REDD+ project in reducing deforestation in the Brazilian Amazon. American Journal of Agricultural Economics, 101, 211–229. DOI: 10.1093/ajae/aay028

SRTM (2020). USGS EROS Archive - Digital Elevation - Shuttle Radar Topography Mission (SRTM) 1 Arc-Second Global. DOI: 10.5066/F7PR7TFT

Stohlgren, T.J., Jarnevich, C.S., Esaias, W.E., & Morisette, J.T. (2011). Bounding species distribution models. Current Zoology, 57, 642–647. DOI: 10.1093/czoolo/57.5.642

Thorn, J.S., Nijman, V., Smith, D., & Nekaris, K. A. (2009). Ecological niche modelling as a technique for assessing threats and setting conservation priorities for Asian slow lorises (Primates: Nycticebus). Diversity and Distributions, 15, 289–298. DOI: 10.1111/j.1472-4642.2008.00535.x

UNEP-WCMC & IUCN (2020). Protected Planet: The World Database on Protected Areas (WDPA)/The Global Database on Protected Areas Management Effectiveness, Cambridge, UK: UNEP-WCMC and IUCN. www.protectedplanet.net. [accessed 5 May 2020].

Wong, M.H., Li, R., Xu, M., & Long, Y. (2013). An integrative approach to assessing the potential impacts of climate change on the Yunnan snub-nosed monkey. Biological Conservation, 158, 401–409. DOI: 10.1016/j.biocon.2012.08.030

Wright, P.C. (1981). The night monkeys, genus Aotus. In Ecology and Behavior of Neotropical Primates (eds F. Coimbra-Filho, R.A. Mittermeier), pp. 211–240. Rio de Janeiro, Academic Brasileira de.

Wright, P.C. (2011). The neotropical primate adaptation to nocturnality: feeding in the night (*Aotus nigriceps and A. azarae*). In: Norconk, M.; Rosenberger, A.L.; Garber, P.A. (Ed.). Adaptive Radiations of Neotropical Primates. Plenum Press, New York. p.369–382. DOI: 10.1007/978-1-4419-8770-9_21

